# Biophysical modeling of neural plasticity induced by transcranial magnetic stimulation

**DOI:** 10.1101/175893

**Authors:** Marcus T. Wilson, Ben D. Fulcher, Park K. Fung, Peter A. Robinson, Alex Fornito, Nigel C. Rogasch

## Abstract

Transcranial magnetic stimulation (TMS) is a widely used noninvasive brain stimulation method capable of inducing plastic reorganisation of cortical circuits in humans. Changes in neural activity following TMS are often attributed to synaptic plasticity (e.g long-term potentiation and depression; LTP/LTD). However, the precise way in which synaptic processes such as LTP/LTD modulate the activity of large populations of neurons, as stimulated en masse by TMS, are unclear. The recent development of biophysically-informed models, which capture the physiological properties of TMS-induced plasticity using mathematics, provide an excellent framework for reconciling synaptic and macroscopic plasticity. In this article, we overview the TMS paradigms used to induce plasticity, and their limitations. We then describe the development of biophysically-based numerical models of the mechanisms underlying LTP/LTD on population-level neuronal activity, and the application of these models to TMS plasticity paradigms, including theta burst and paired associative stimulation. Finally, we outline how modeling can complement experiment to improve mechanistic understandings and optimize outcomes of TMS-induced plasticity.

**Abbreviations:**

TMS: transcranial magnetic stimulation
LTP: long-term potentiation
LTD: long-term depression
rTMS: repetitive TMS
TBS: theta burst stimulation
PAS: paired-associative stimulation
MEP: motor-evoked potential
cTBS: continuous TBS
iTBS: intermittent TBS
NMDA: n-methyl-d-aspartate
CaDP: calcium dependent plasticity
EEG: electroencephalography
MRI: magnetic resonance imaging
STDP: spike timing dependent plasticity
BCM: Bienenstock-Cooper-Munro
GABA: gamma-aminobutyric-acid
ISI: inter-stimulus interval

## 1. Introduction

Transcranial magnetic stimulation (TMS) is a powerful tool for studying human brain function (Hallett, 2007). Using the principles of electromagnetic induction, TMS can noninvasively depolarize cortical neurons across the skull and scalp (Rothwell, 1997). When applied using certain patterns of stimulation, TMS can induce plastic increases or decreases in cortical excitability which outlast the period of stimulation, reminiscent of long-term potentiation/depression (LTP/LTD)-like mechanisms observed in vitro (Cooke and Bliss, 2006). The ability of TMS to alter neural activity has resulted in a wide range of uses, from studying the mechanisms of plasticity in humans (Ziemann et al., 2008), to inferring the functional roles of brain regions in behavior (Pascual-Leone et al., 2000), to providing clinical treatments (Lefaucheur et al., 2014). However, the field is facing several challenges, including large interindividual variability in response to TMS (Ridding and Ziemann, 2010), difficulty in optimizing protocols due to nonlinear relationships between stimulation parameters and outcomes (Gamboa et al., 2010), and problems translating the detailed knowledge gained from the motor system to non-motor regions which often lack clearly observable output that can be used to measure the effects of stimulation. (Parkin et al., 2015). At the centre of these challenges is a poor understanding of the cellular and molecular mechanisms underlying TMS-induced plasticity (Müller-Dahlhaus and Vlachos, 2013), which makes it difficult to interpret outcomes at the level of neural populations as measured in human experiments. As such, a general theory that explains how TMS induces plasticity across multiple brain scales and regions is urgently required.

Our aim here is to overview how biophysically-informed modeling approaches can be applied to better understand TMS-induced plasticity thus addressing the challenges outlined above. These models aim to explain and predict data in terms of underlying physiological mechanisms. The central idea is to capture the relevant physiological mechanisms using mathematics, where the variables and parameters of the model equations reflect physiological properties such as receptor conductances, membrane potentials, or firing rates. We begin by briefly overviewing the TMS paradigms used to induce plasticity in humans, the proposed mechanisms underlying these protocols, and the practical and theoretical factors limiting the use and interpretability of these paradigms. We then introduce how biophysically-based models, including neural field models, have been used to describe neural plasticity mechanisms. Finally, we overview the application of modeling to TMS plasticity paradigms, discuss how such models can inform the optimization of plasticity-inducing paradigms for TMS, and outline future research directions to further refine our understanding of the interactions between TMS and plasticity across different brain regions. Modeling will play a key role in resolving many of the seemingly inconsistent and unexpected findings in the field of TMS research and we argue for greater integration of modeling and experimentation to guide the future of the field.

## 2. Plasticity-inducing TMS paradigms

To illustrate the need for models of TMS induced plasticity, we first introduce the common TMS paradigms used to induce plasticity in humans, the proposed molecular mechanisms underlying these paradigms, and the challenges facing the field of TMS. The three most commonly used plasticity-inducing TMS paradigms are repetitive TMS (rTMS), theta burst stimulation (TBS), and paired associative stimulation (PAS) (Ziemann et al., 2008). Most research to assess the capacity of these paradigms to induce plasticity has occurred in the motor system due to the ease of measuring motor outputs. For instance, single TMS pulses given to the motor cortex at suprathreshold intensities result in compound action potentials in the muscle targeted by the stimulated cortical region. This TMS-evoked muscle activity, termed a motor-evoked potential (MEP), can be easily measured using surface electromyography. The amplitude of the MEP is influenced by both cortical and spinal excitability, but is often used to indirectly infer changes in cortical excitability induced following TMS plasticity protocols (Di Lazzaro et al., 2008).

All three TMS plasticity protocols listed above can either increase or decrease MEP amplitude for ∼15-30 minutes following stimulation. However, the stimulation protocols through which these excitability changes are achieved differ between paradigms. rTMS involves administering repeated uniformly-spaced pulses of TMS, with the after-effects depending predominantly on the frequency of stimulation. When applied at lower frequencies (e.g., at 1 pulse per second), rTMS generally results in a decrease in MEP amplitude, suggesting reduced cortical excitability. Conversely, higher rTMS frequencies (> 5 Hz) usually result in increased MEP amplitudes (Fitzgerald et al., 2006).

In TBS, there are two frequencies of stimulation - a slower interburst ‘theta’ frequency (usually 5 Hz, meaning successive bursts of pulses are separated by 200 ms), and also a faster intraburst ‘gamma’ frequency (usually 50 Hz, meaning pulses within a burst are separated by 20 ms). The number of pulses in a burst is also variable (though often set to 3); likewise the total number of pulses in a protocol, often set to 600, can be varied. Protocols can be applied continuously (cTBS) or intermittently (iTBS); in the latter pulses are delivered for a given period (the ‘ON’ time, often 2 s) and then are absent for a given period (the ‘OFF’ time, often 8 s). The excitability effects of TBS depend on the pattern of stimulation. It is commonly assumed that cTBS decreases MEP amplitude, whereas iTBS increases MEP amplitude (Huang et al., 2005).

PAS uses a different approach to rTMS and TBS, and is based on the principles of ‘Hebbian’ or spike timing dependent plasticity (Stefan et al., 2000). During PAS, single TMS pulses are delivered to the motor cortex to coincide with sensory inputs resulting from electrical stimulation of a peripheral nerve (Stefan et al., 2000). The changes in cortical excitability resulting from PAS are dependent on the relative timing of the two stimuli. For example, at an interstimulus interval of ∼25 ms (PAS_25_), the sensory input, which has a conduction time to the cortex of ∼20 ms, and the TMS pulse arrive at a similar time, yielding increased MEP amplitudes. Conversely, at an interstimulus interval of ∼10 ms (PAS_10_), the sensory input arrives well after the TMS pulse, resulting in reduced MEP amplitudes (Wolters et al., 2003).

In addition to the frequency and pattern of stimulation, other parameters such as stimulation intensity, number of pulses, and brain state also influence the outcome of each paradigm (Pell et al., 2011). Furthermore, the generally accepted ‘rules’ that high frequency rTMS, iTBS, and PAS_25_ increase excitability and low frequency rTMS, cTBS, and PAS_10_ decrease excitability do not hold in all people (see *Challenges facing TMS plasticity paradigms*) (Hamada et al., 2013; Maeda et al., 2000; Mϋller-Dahlhaus et al., 2008).

## 3. Proposed mechanisms underlying TMS-induced plasticity

Each TMS plasticity protocol is motivated by electrical stimulation paradigms used to induce LTP/LTD across single synapses in hippocampal slice studies in animals (Cooke and Bliss, 2006; Raymond, 2007). In these studies, LTP/LTD primarily alters synaptic efficacy by increasing or decreasing the conductance and number of AMPA receptors at the postsynaptic neuron, with presynaptic mechanisms such as increased or decreased neurotransmitter release also playing a role (Dan and Poo, 2004; Malenka and Bear, 2004). The direction of synaptic plasticity depends on postsynaptic intracellular calcium concentrations (hence this is also known as calcium dependent plasticity; CaDP). TMS activates different neural populations depending on the intensity of the stimulation and the geometry of the coil, leading to action potentials. In the theory of CaDP, depolarization of a postsynaptic neuron leads to calcium influx to the postsynaptic dendritic spine via glutamatergic, voltage-gated NMDA receptors (Dan and Poo, 2004). From then on, complex chains of protein kinases and phosphatases may be activated, so that low calcium concentrations result in LTD, and higher concentrations result in LTP (Nishiyama et al., 2000; Shouval et al., 2002). The direction and magnitude of synaptic plasticity is further governed by the history of synaptic activity, a process known as metaplasticity (Abraham, 2008). Several mechanisms have been suggested to account for metaplasticity, including a sliding plasticity threshold mediated by changes in NMDA receptor conductance (Bienenstock et al., 1982; Shouval et al., 2002).

Changes in MEP amplitude from all three TMS protocols are broadly consistent with changes in synaptic efficacy following LTP/LTD-like mechanisms (Hoogendam et al., 2010). First, the changes in excitability induced by TMS paradigms last beyond the period of stimulation for short periods (∼30 min), consistent with LTP/LTD (Huang et al., 2005; Peinemann et al., 2004; Stefan et al., 2000). Second, the effects of TMS paradigms are blocked or reversed by NMDA receptor (Fitzgerald et al., 2005; Huang et al., 2007; Stefan et al., 2002) and calcium channel antagonists (Wankerl et al., 2010; Weise et al., 2016). Third, the outcomes of TMS plasticity paradigms are dependent on the history of cortical activation (e.g., past plasticity induced by another TMS protocol or motor learning), consistent with metaplasticity (Iyer et al., 2003; Müller et al., 2007; Todd et al., 2009; Ziemann et al., 2004). In addition to synaptic plasticity, other nonsynaptic mechanisms may also contribute to changes in MEP amplitude, such as changes in membrane excitability, biochemistry, or gene expression (Pell et al., 2011; Tang et al., 2015). However, the contributions of these mechanisms to TMS plasticity are yet to be fully explored.

## 4. Challenges facing TMS plasticity paradigms

Despite the widespread use of TMS, several challenges currently limit the design and interpretability of TMS-induced plasticity experiments. First, it is unclear how the plasticity mechanisms that likely underlie the effects of TMS, such as LTP/LTD, scale from the microscale/synapse level in animal studies to the macroscale/brain region level in human studies (Müller-Dahlhaus and Vlachos, 2013). For example, in animal studies, plasticity is induced following selective stimulation of one or several presynaptic, usually excitatory, neurons (Raymond, 2007). In contrast, TMS depolarizes large populations of both excitatory and inhibitory neurons at the pre- and postsynaptic level (Di Lazzaro and Ziemann, 2013), meaning that TMS may induce plasticity in both excitatory and inhibitory neural populations (Chung et al., 2016; McAllister et al., 2009). Methods that bridge micro-, meso-, and macroscale plasticity and can study multiple neural populations are therefore needed to better establish the cellular and molecular mechanisms of TMS-induced plasticity. The development of animal (Funke and Benali, 2011) and hippocampal cell culture (Vlachos et al., 2012) preparations for studying TMS have allowed more precise assessments of the specific neural populations altered by stimulation, and the molecular mechanisms underlying these changes. However, it is difficult to reproduce equivalent spatial distributions of electric fields across scales, since TMS typically stimulates large sections of the brain, if not the whole body, of smaller animals like rodents (Tang et al., 2015).

The lack of a strong mechanistic understanding of TMS-induced plasticity at the level of neural populations likely contributes to the second major challenge for the field: the large interindividual variability in response to TMS plasticity paradigms. Recent studies have demonstrated that only ∼50% of participants show the expected plasticity effects following a particular stimulation protocol (Hamada et al., 2013; Hinder et al., 2014; Lόpez-Alonso et al., 2014; Maeda et al., 2000; Müller-Dahlhaus et al., 2008). For example, only 50% of the participants in (Hamada et "https://paperpile.com/c/tL6ALY/19rB" al., 2013) showed the expected increase in MEP amplitude after iTBS in the motor cortex. Such variability extends to behavioral (Nicolo et al., 2015) and clinical outcomes (Lefaucheur et al., 2014) of TMS, and severely limits the reliability and hence usefulness of TMS as an experimental and clinical tool.

The reasons for this interindividual variability remain unclear but several factors likely contribute, including the state of the cortex (e.g. engaged in a conscious task, following a muscle contraction), age, sex, genetics, neurotransmitter and receptor variation, connectivity of the stimulated region, position of the coil relative to the target population, geometry of the head, and the neural population stimulated by TMS (see Ridding and Ziemann, 2010). Another factor that is likely key, but has received surprisingly little attention, is the choice of stimulation protocol and parameters (Goldsworthy et al., 2012). Taking TBS as an example, the number of possible stimulation protocols is enormous, with freedom to independently set: (i) the frequency of burst stimulation; (ii) the burst interval; (iii) the number of pulses within a burst; (iv) the number of pulses in a session; and (v) the intensity of stimulation (pulse amplitude and duration). However, nearly all TBS studies have used 50 Hz bursts at 5 Hz intervals for 600 pulses at 80% of active motor threshold (Chung et al., 2016). While the repeated use of these ‘standard’ parameters likely reflects a desire to use protocols that have been shown to induce plasticity and have demonstrated safety, the few studies that have altered either frequency (Goldsworthy et al., 2012), pulse number (Gamboa et al., 2010; Gentner et al., 2008), or intensity (Doeltgen and Ridding, 2011) have observed nonlinear relationships between parameter changes and plasticity. Such findings highlight the difficulty of searching the parameter space experimentally, without an underlying theory to guide parameter selection. Given the large interindividual variability in plasticity following TBS (Hamada et al., 2013), it is also possible that the optimal frequency of stimulation might depend on factors specific to individual participants or certain brain regions, such as the resonant frequency of ongoing neural oscillations (Klimesch et al., 2003) or strengths of connections between regions (Sethi et al., 2017). Methods that are able to guide parameter selection are thus needed to reduce the dimensionality of this problem.

Finally, it is unclear how results from TMS studies in the motor cortex translate to non-motor regions which have no output equivalent to the MEP. The combination of rTMS with neuroimaging methods such as electroencephalography (EEG) (Chung et al., 2015; Rogasch and Fitzgerald, 2013; Thut and Pascual-Leone, 2010) and magnetic resonance imaging (MRI) (Sale et al., 2015) has helped to address this issue, however the relationship between EEG activity, hemodynamics measured with functional MRI, and MEPs remains unresolved. Interestingly, neuroimaging studies have demonstrated that TMS not only induces changes in the stimulated region, but also across extended regions of the brain (Cocchi et al., 2015). Furthermore, identical stimulation to different regions can result in opposite outcomes, with stimulation increasing functional connectivity at one region, but decreasing functional connectivity at another (Cocchi et al., 2016; Eldaief et al., 2011). In order to unify the disparate results obtained from different brain regions using different measurement modalities, we urgently require a framework that structures our results and understanding in terms of putative physiological mechanisms.

## 5. Mathematical modeling and neural plasticity

Biophysically-based models aim to explain and predict data in terms of underlying physiological mechanisms. Computers are used to simulate the models of the neurophysiology, treating them as forward models to yield predictions of the data that can then be measured experimentally. Conversely, given experimental data, the models can be inverted to infer model parameters (which putatively correspond directly to physiological quantities) that best explain the data. In this way, mathematical models of the brain act as a bridge between experimental data, which are indirect measurements of the brain, and their generative, unobserved neurophysiological mechanisms.

Mathematical models of the brain can be formulated at multiple levels of description, from the microscopic scale of single neurons, through to the interplay of macroscopic neural elements to produce complex patterns of whole brain dynamics (Breakspear, 2017; Deco et al., 2008). The appropriate level of description to include in any model depends on the question of interest and scale of analysis. Including too much complexity may lead to an under-constrained model that could fit any phenomenon, but simplifying too much may obscure phenomena of interest. In humans, for example, neural activity is most commonly measured at the macroscale, e.g., using MRI or EEG, with data reflecting the combined activity of extended populations of neurons. For data at this macroscopic scale, rather than explicitly simulating an incredibly complicated and under-constrained model of the activity of billions of individual neurons, it is more appropriate to reduce large populations of spiking neurons to their collective properties (e.g., the mean firing rate of neurons in each population). This approach to macroscopic brain modeling is appropriate for the electrical stimulation of the brain by externally applied fields such as TMS, which stimulates bulk populations of neurons.

*Mean field models* are a class of models appropriate for describing the macroscopic scale of large populations of neuron distributed across space. Spatially distributed populations of neurons form the basic elements of these models, with equations governing how overall properties of these populations evolve through local dynamics and interactions with other populations, for example (Deco et al., 2008). *Neural field models* take into account the continuous spatial extent of the brain and can, for example, capture propagation of waves across the cortex (Jirsa and Haken, 1996; Robinson et al., 2001). A simplification yields *neural mass models*, where each neural population is represented as a single number (the mean of the neurons in that population). Neural masses at different locations can be linked together, but this must be done in a way that is robust to finer discretization. Model equations and their parameters encapsulate interpretable (and often directly or indirectly biologically measurable) mechanisms considered appropriate for capturing the phenomenon of interest, such as interaction parameters that control the interplay of different populations (such as inhibition by inhibitory interneurons). Once constructed, models can be analyzed mathematically, or simulated numerically with a computer to fit experimental data and generate predictions to guide future experiments (Bojak and Liley, 2010; Deco et al., 2008; Jirsa and Haken, 1996; Nunez, 1974; Robinson, 2005; Robinson et al., 2005, 1997; Steyn-Ross et al., 2009).

Mean field models have proved successful in interpreting diverse biophysical phenomena, including the sleep-wake transition (Phillips and Robinson, 2007; Steyn-Ross et al., 2005), EEG spectra from 0.1 - 100 Hz (Robinson et al., 2001; Wright and Liley, 1996), evoked potentials such as sensory event-related potentials (Rennie et al., 2002), epileptic seizures (Kramer et al., 2005; Robinson et al., 2002), low frequency spontaneous fluctuations of the blood oxygenation level-dependent signal recorded with MRI (Steyn-Ross et al., 2009), and visual hallucinations under migraine (Henke et al., 2014a, 2014b). Artificially altered brain states such as general anesthesia have also been modeled successfully (Bojak and Liley, 2005; Wilson et al., 2006).

Figure 1 shows some examples of coupled neuronal population models, including: (a) a single population of cortical excitatory neurons (Robinson, 2011), (b) a coupled population of excitatory and inhibitory cortical neurons (Fung et al., 2013), and (c) a corticothalamic model including excitatory and inhibitory cortical neurons with populations of thalamic reticular and thalamocortical relay neurons. All can be stimulated with TMS protocols by adding in a ‘TMS’ population with a given rate of firing as a function of time, as shown here, or with other forms of stimulation.

**Figure 1.**
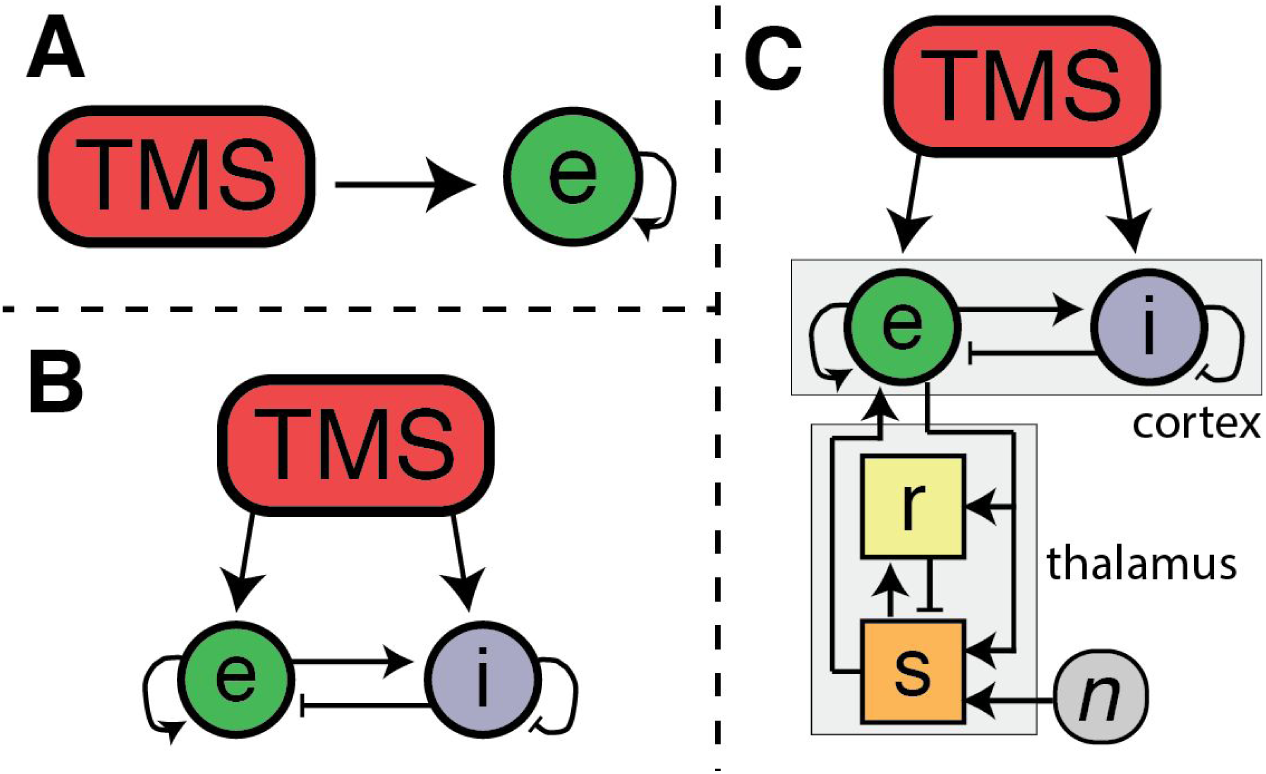
*Schematic representations of three examples of neuronal populations and their interactions that can be numerically simulated using biophysical models.* ***A*** *A single population of cortical excitatory neurons, e, driven by an external TMS stimulus.* ***B*** *Coupled excitatory, e, and inhibitory, i, cortical neuronal populations.* ***C*** *Coupled cortical (excitatory, e, and inhibitory, i, neurons) and thalamic (reticular, r, and thalamocortical relay, s) neuronal populations, with a separate subcortical noise term n which represents sensory input. The progression A - C allows phenomena of different complexities to be captured; for example cortical excitatory-excitatory plasticity can be investigated through A; general anesthesia, which involves lengthening inhibitory post-synaptic potentials, requires consideration of inhibitory cortical neurons as in B; the alpha resonance of the EEG requires thalamic processes to be added, as in C. In all three cases populations can coupled to themselves, meaning the future activity of a population is directly influenced by its current activity.*

We now consider some details of mean field models. Although different authors have used different formulations, all such models use mathematical equations to describe the dynamics of populations of neurons and the interactions between them. We used the scheme of Fig. 1B as an example. Figure 2A shows the coupled populations in more detail, indicating some of the key biophysical parameters of the model, including coupling strengths between populations, *v*_*jk*_ (which specify the strength of excitation or inhibition between populations), and timescales, *τ*_*j*_ (which set the intrinsic timescale of activity for a population). Setting these parameters specifies the model; for example, setting *v*_*ei*_ < 0 specifies inhibition from population *i*→*e*, setting *v*_*ie*_ > 0 specifies excitation from population *e*→*i*, and increasing *τ*_*e*_ increases the response timescale for the excitatory population, e. Neuronal populations are represented by the mean cell body potential across neurons of the population, *V*_*i*_, which can simulated from the parametrized set of interactions and time constants. This mean membrane potential can be related to the mean firing rate through a parametrized sigmoidal function (Fig. 2B) (Freeman, 1975). Different cortical or subcortical regions can be considered as discrete, but connected, neural populations. To describe interactions between these neural populations, specific parameters quantify the strength of synaptic coupling, and the direction, magnitude, and time course of synaptic input to each population (Fig. 2C). Plasticity can be incorporated phenomenologically through an adaptation of a pairwise spike timing dependent plasticity (STDP) window (Bi and Poo, 2001) for neuronal populations. STDP requires adaptation to capture spike triplet interactions (such as two presynaptic spikes paired with one postsynaptic spike) which are important when firing rates are high and spikes are close together (Pfister and Gerstner, 2006). Since several hundred presynaptic spikes typically occur in the integration time for a cortical neuron to produce a postsynaptic spike, population based approaches to plasticity are particularly suitable (Deco et al., 2008).

**Figure 2.**
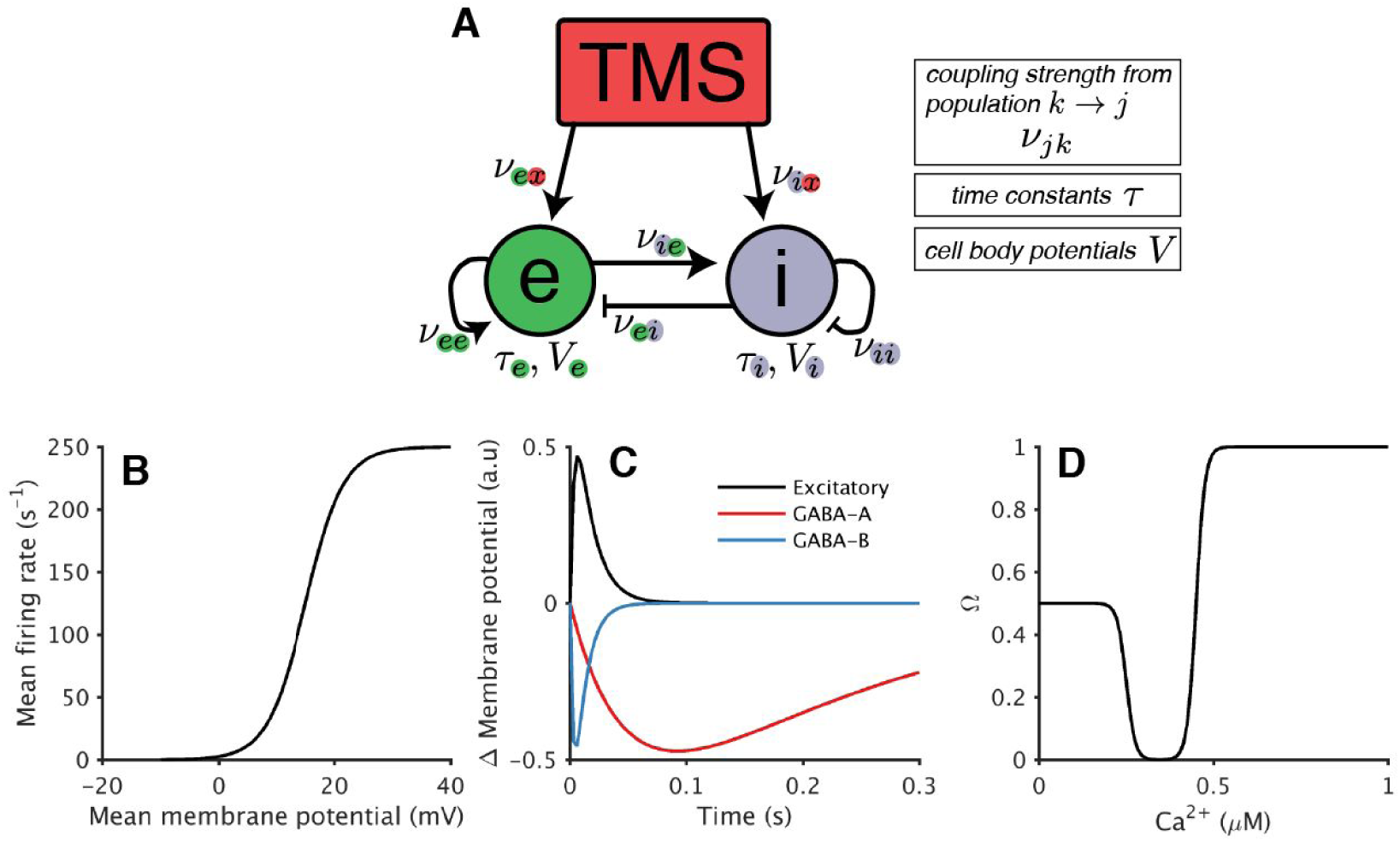
*Three examples of how physiological relationships can be approximated in models of macroscale brain activity. (A) Populations of the model of Fig. 1B, annotated with some relevant parameters, including coupling strengths between populations, v*_*jk*_, *timescales, τ*_*j*_, *and cell body potentials, V*_*j*_. *(B) The relationship between mean cell body (membrane) potential V*_*j*_ *and mean firing rate within a neural population can be described with a sigmoid function (Freeman, 1975).(C) Changes in membrane potential V*_*e*_ *across time resulting from a single input of an excitatory population synapsing onto dendritic excitatory receptors (black line), and an inhibitory population synapsing onto dendritic GABA*_*A*_ *(red) and GABA*_*B*_ *(blue) receptors. The amplitude of the change in membrane potential has been scaled to illustrate differences in peak change between receptor types. The GABAA and GABAB curves describe the dynamics of the inhibitory connections *i*→*e* and *i*→*i* of part (A); the excitatory curve describes the dynamics of the excitatory connections *e*→*e* and *e*→*i*. (D) The Ω synaptic plasticity function codes the direction of change of synaptic strengths *v*_*ee*_. Neural firing results in calcium influx to the cell, the concentration of which determines whether synaptic strength decreases (Ω < 0.5; LTD) or increases (Ω > 0.5; LTP).*

More detailed approaches have been developed to include calcium-dependent plasticity (CaDP) (Graupner and Brunel, 2010). CaDP theory (Shouval et al., 2002) describes cellular calcium dynamics and its effects on synaptic strength. The calcium concentration levels resulting in depression or potentiation are described through the Ω function (Fig. 2D): Ω<0.5 gives LTD; Ω>0.5 gives LTP. CaDP theory has been incorporated into mean field models (Fung and Robinson, 2014),(Huang et al., 2011). Calcium influx into postsynaptic dendritic spines through NMDA receptors is dependent on both glutamate release due to presynaptic activity and postsynaptic voltage, so CaDP provides a microscopic link between the activity of pre- and postsynaptic populations of cells. The theory has been further expanded to include a Bienenstock-Cooper-Munro (BCM) metaplasticity scheme (Fung and Robinson, 2014). Here, activity-dependent changes in NMDA receptor calcium conductance are assumed to underlie metaplasticity; previously high plasticity induction reduces calcium conductance and favors LTD, while previously low induction increases conductance favoring LTP (Bienenstock et al., 1982). As such, the metaplasticity scheme operates as an activity-dependent sliding window, similar to the BCM postulate, meaning that the requirements for inducing plasticity depend on the history of plasticity induction.

## 6. Modeling of TMS plasticity paradigms

The models considered above provide useful tools for understanding the effects of TMS on plasticity. We now consider in more detail how such models have been applied, specifically to rTMS, TBS, and then PAS. The model results are compared to experiments in each case. We then show how modeling can provide insights to guide experimental design and interpretation of the results of TMS protocols.

The rTMS paradigm has been studied with models of STDP (Fung et al., 2013) and CaDP (Fung and Robinson, 2013). In these models, changes in synaptic coupling between excitatory populations were used to estimate the TMS-induced change in cortical excitability, measured experimentally as MEPs. Results obtained using a CaDP model are summarized in Fig. 3A for one-second pulse trains. Low frequencies (<8 Hz) resulted in little change in synaptic weight, higher frequencies (8-14 Hz) resulted in LTD, while yet higher frequencies (>14 Hz) gave very strong LTP. Human rTMS studies generally show LTD-like effects at 1 Hz and LTP-like effects above 5 Hz. The model has reproduced the general pattern of LTD at low and LTP at high frequencies, but the frequency ranges are not correct (Fitzgerald et al., 2006). It remains unclear whether using stimulation paradigms that more closely resemble those used in human experiments (e.g., 5 Hz stimulation with 5 s on and 25 s off over 1500 pulses), or including a metaplasticity scheme (Fung and Robinson, 2014), would alter these results.

**Figure 3.**
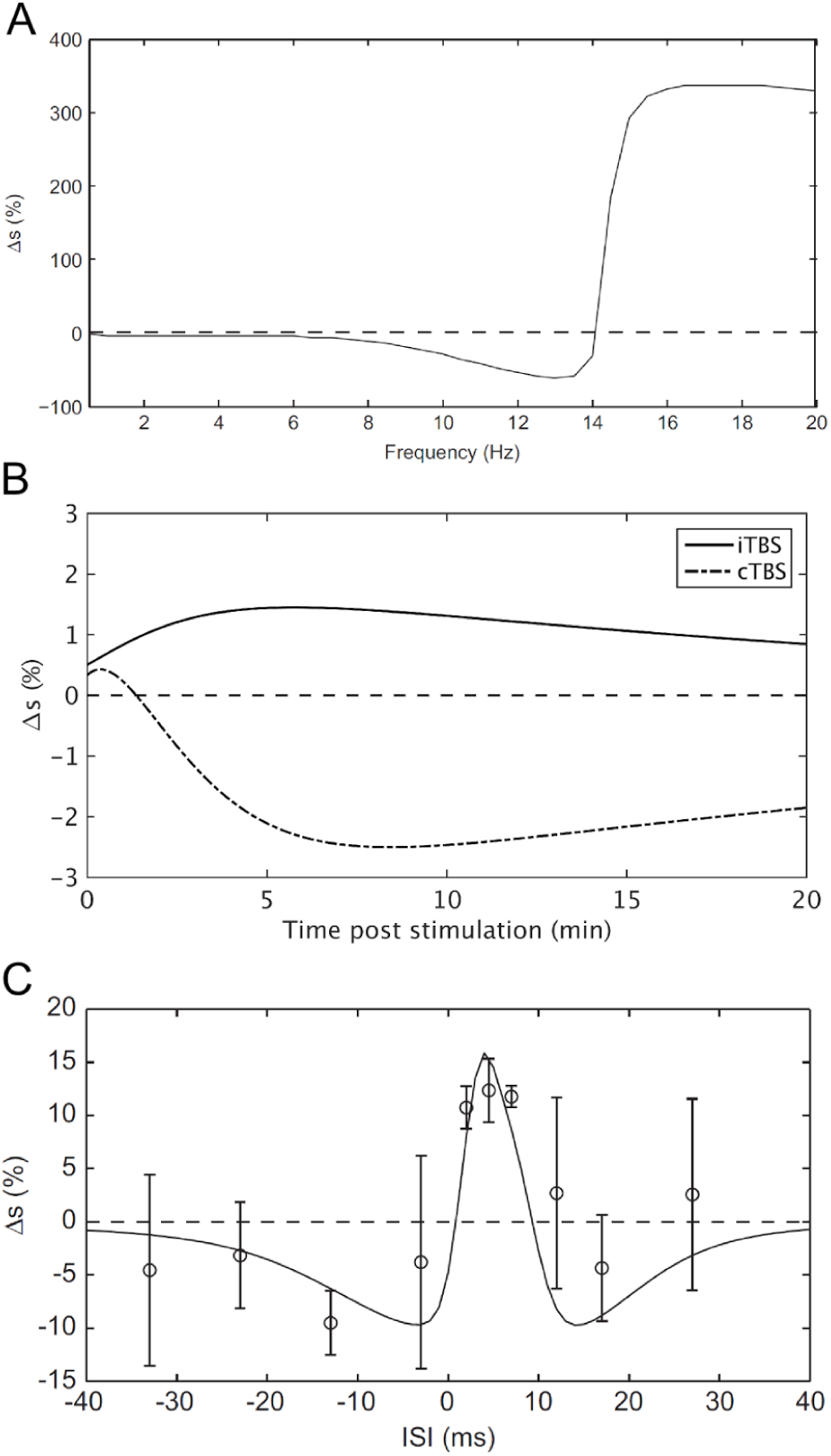
*Plasticity effects of different TMS paradigms predicted by a single excitatory population CaDP model. A The effect of rTMS frequency on change in synaptic coupling between excitatory populations (Δs). B The effect of iTBS and cTBS on excitatory synaptic coupling. C The effect of interstimulus interval during PAS on the change in excitatory synaptic coupling (solid line). The circles and error bars represent changes in MEP amplitude from human participants (Wolters et al., 2003). A and C are taken from (Fung and Robinson, 2013) with permission.*

Theta burst stimulation protocols were considered from a theoretical perspective by Huang et al. (Huang et al., 2011). They proposed a model of plasticity based on ‘facilitatory’ and ‘inhibitory’ agents. Both were driven by postsynaptic calcium concentration; however, the former depended upon the rate of increase of calcium while the latter depended on the *accumulated* concentration of calcium. Many of the canonical TBS results could be explained in this manner; cTBS (600 pulses) was ultimately depressive due to the accumulation of postsynaptic calcium, but iTBS was potentiating since the breaks in pulses allowed for calcium levels to decay between bursts. Other phenomena were also captured – for example a 300 pulse cTBS protocol without prior muscle contraction did not produce enough calcium for the inhibitory effects to dominate and was hence potentiating, but with prior muscle contraction the calcium level was driven to higher levels and the protocol became potentiating. This last result, the effect of previous activity, is an example of *metaplasticity*. However, the model failed to capture other notable TBS results, for example the switch in plasticity direction for cTBS (from LTD to LTP) and iTBS (from LTP to LTD) when a 600 pulse protocol was extended to 1200 pulses (Gamboa et al., 2010).

Neural masses have also been used to model the TBS paradigm (Fung and Robinson, 2014). CaDP, with the addition of metaplasticity in a BCM scheme, was included to show that a biophysically detailed plasticity model can reproduce the effects of TBS. A single excitatory population of cells was modeled (as in Fig. 1A), and the plasticity response (calculated as the change in synaptic coupling between excitatory populations, Δs) for the canonical cTBS and iTBS protocols (Huang et al., 2005) was simulated. The model predicted the canonical LTD and LTP for cTBS and iTBS, respectively (Fig. 3B). Of particular importance was the predicted dependence of the plasticity response on the number of pulses in the protocol. Either doubling or halving the number of pulses reversed the TBS outcome, with cTBS resulting in LTP and iTBS resulting in LTD, also consistent with experimental findings (Gamboa et al., 2010; Gentner et al., 2008). Furthermore, reducing TMS activation of the excitatory population also reversed the excitability change following both iTBS and cTBS. This pattern is consistent with the finding in humans that differences in activation of excitatory and interneuron populations by the TMS pulse (e.g., differences in MEP latency following stimulation with anterior-posterior current flow) are tightly correlated with the resulting TBS outcome (Hamada et al., 2013). The increase/decrease in excitability following iTBS/cTBS has also been replicated using extended CaDP (Wilson et al., 2016) and STDP models (Wilson et al., 2014) incorporating both an excitatory and inhibitory population of cells (as in Fig. 1B), and including realistic synaptic response times for both GABAA and GABAB receptors as in Fig. 2C. Results from the STDP model were close to predictions from a model including only CaDP, indicating realistic STDP windows can be recovered from CaDP predictions (Wilson et al., 2016).

PAS has been modeled using CaDP (Fung and Robinson, 2013). The plasticity at different interstimulus intervals (ISI) was calculated for a single population of excitatory cells (Fig. 1A). This model predicted depression for ISI between approximately -20 to 0 ms (negative times meaning that the TMS pulse occurred before the nerve stimulus arrived), potentiation for ISI between approximately 0 and 10 ms, and a second small depressive window for longer ISI, from about 10 to 30 ms, roughly in agreement with experiment (Bi and Poo, 2001; Graupner and Brunel, 2010). The predictions are shown in Fig. 3C. Taken together, these findings demonstrate that mean field models incorporating CaDP are capable of reproducing the main patterns of excitability change following all three major TMS plasticity paradigms. These results provide evidence for CaDP being a key mechanism of macroscale TMS-induced plasticity.

## 7. Understanding interindividual variability and model-based optimization of stimulation parameters

One of the major factors limiting the practical use of TMS plasticity paradigms is the large interindividual variability in response. Modeling approaches, which allow experimental results to be interpreted in terms of inferred neurophysiological parameters, may be able to provide new insights into the reasons behind these broad interindividual responses to TMS and thereby help inform the design of future experiments.

As an example, the CaDP model of TBS with metaplasticity has been used to simulate synaptic strength over time following iTBS and cTBS protocols, as shown in Fig. 4. Results show that the change in synaptic strength following a protocol depends upon the duration of the protocol. This dependency arises in the model as a result of the interplay between calcium concentration, plasticity signaling, and metaplasticity, giving rise to an oscillatory behavior, with the direction of change in synaptic coupling dependent on the concentration at the end of stimulation (Fung and Robinson, 2014). The implication of this finding is that *both* iTBS and cTBS are capable of producing increases *and* decreases in cortical excitability depending on the length of stimulation. Furthermore, the phase of the calcium oscillation during stimulation is shifted by altering the level of TMS activation on the excitatory population (Fung and Robinson, 2014). As such, the number of pulses required for an increase/decrease in excitability following iTBS/cTBS may differ between individuals depending on how TMS interacts with excitatory cortical populations. This may be due to the differences in structure of brain convolutions or genetic differences in physiology between individuals, with the same applied field strength and pattern activating slightly different groups of neurons. These results suggest that differences in the response to TBS between individuals can be minimized by adequately adjusting either the number of pulses given, or the TMS intensity for the individual. It is possible that by identifying and understanding the elements of physiology that most affect TMS response, we may be able to build personalized models, and use these models to tailor clinical treatments.

**Figure 4.**
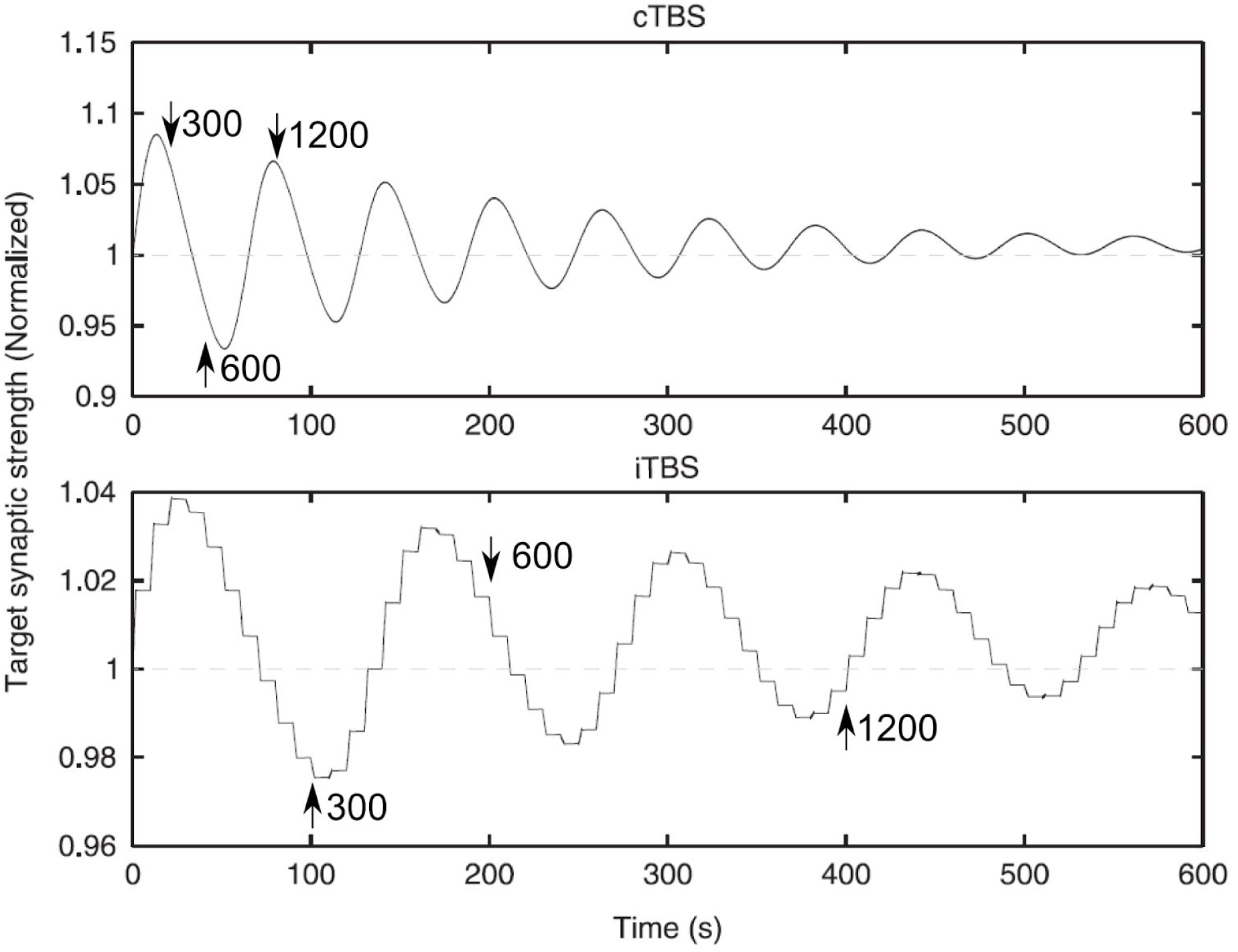
*Predicted change in synaptic strength as a result of oscillations in calcium concentration during cTBS (top) and iTBS (bottom). Vertical axes indicate the synaptic strength attained (relative to the initial strength) if the stimulation protocol is ceased at the point in time indicated on the horizontal axis. Arrows indicate the change in excitability for different numbers of pulses. Following the standard 600 -pulse protocol, the canonical decrease/increase in excitability with cTBS/iTBS is predicted by the model. However, for 300 pulses and 1200 pulses, the model predicts a reversal of the above pattern. Figure taken from (Fung and Robinson, 2014) with permission.*

Additionally, modeling can be used to explore the vast stimulation parameter space to inform optimization of TMS paradigms. A benefit of capturing the relevant physiological dynamics and interactions in a model is that alterations in parameters can be simulated numerically, without the need to perform costly experiments. This is of particular use for protocols like TBS, where the vast parameter space defining myriad combinations of stimulation timing and strength remains mostly unexplored, but can be simulated straightforwardly (Wilson et al., 2016, 2014).

Figure 5 shows the predicted change in MEP strength after 600 pulses of close-to-threshold cTBS (part A), and iTBS (2 s ‘ON’ time, 8 s ‘OFF’ time, part B) for a wide range of different interburst (theta) stimulation frequencies, intraburst (gamma) frequencies, and pulses-per-burst applied to the neuronal model of (Wilson et al., 2016) (Fig. 1B). The canonical cTBS and iTBS protocols (Huang et al., 2005) are part of this set; they are shown by the crosses in Fig. 5. It is clear that an increase in intraburst stimulation frequency generally leads to an increase in potentiation. Likewise, higher numbers of pulses per burst favor LTP over LTD. The effect of interburst (theta) frequency is more subtle; for cTBS at lower theta frequencies increasing stimulation rate favors LTD over LTP, but at high theta frequencies and for iTBS the potentiation is not greatly dependent on stimulation rate.

**Figure 5.**
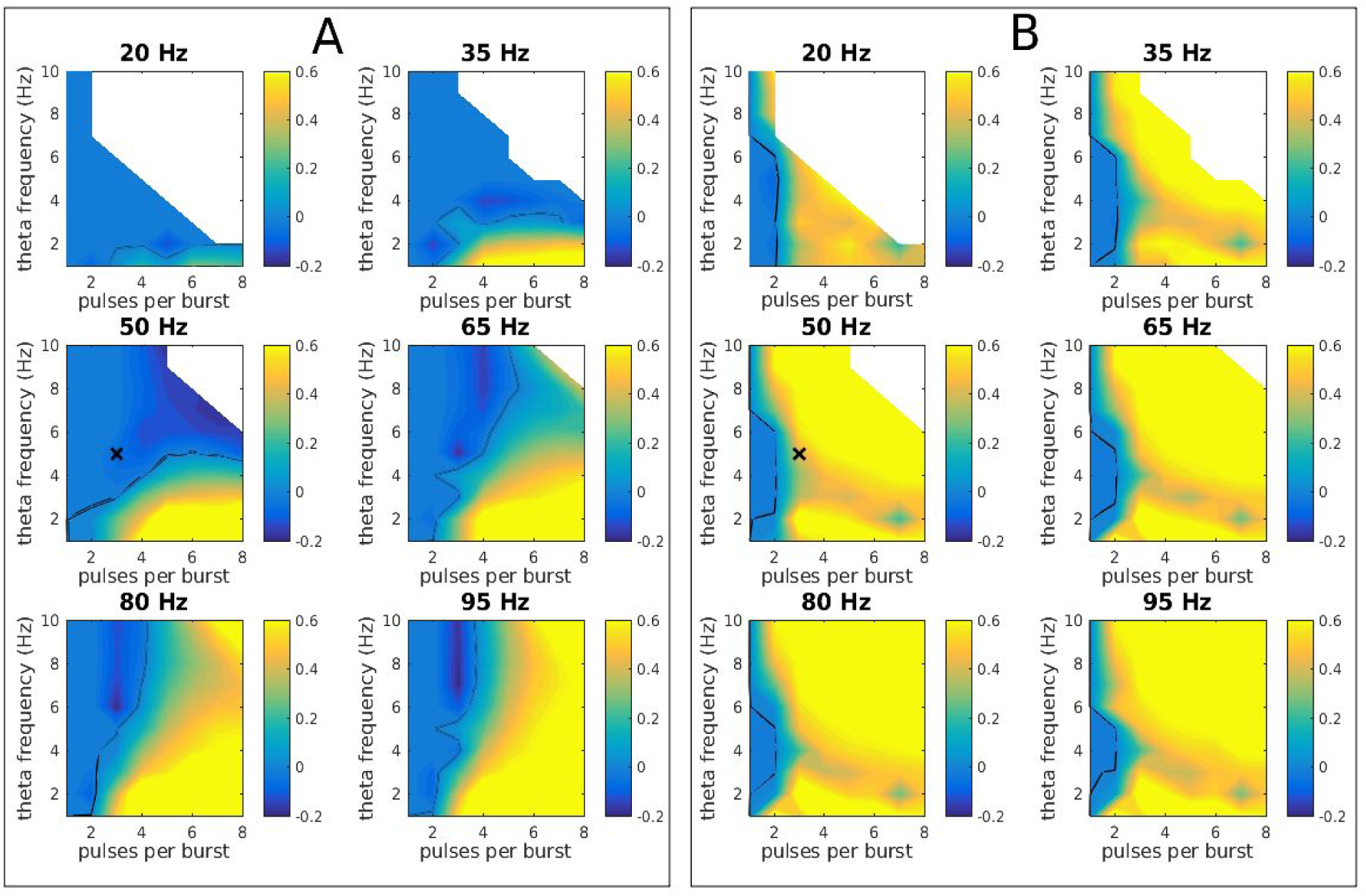
*Estimated change in excitatory-excitatory synaptic strength (CaDP with BCM metaplasticity; coupled excitatory and inhibitory populations) for continuous bursting (A) and intermittent bursting (B) protocols. Parts (A) and (B) both show the relative change in synaptic coupling (0 = no change, positive values shown as green-yellow = LTP, negative values shown as blue = LTD) immediately after 600 pulses of close-to-threshold TBS versus three variables:1. The interburst (theta) stimulating frequency (1 - 10 Hz, on the vertical axis); 2. The number of pulses in a burst (1 - 8, on the horizontal axis); and 3. The intraburst stimulation frequency (20 - 95 Hz, indicated above each set of axes). The solid black contours denote the boundary between LTD and LTP. The ‘X’ symbols in parts (A) and (B) denote respectively the cTBS and iTBS protocols of (Huang et al., 2005).*

The results demonstrate the ease with which constrained models of candidate neurophysiological mechanisms can generate predictions about the results of new experiments, in this case helping the experimenter to understand the impact of stimulation timing parameters for TBS. The impact of such predictions could be important for selecting optimal stimulation settings for addressing a given scientific question, or to obtaining the maximal treatment response; issues that cannot currently be addressed systematically from first principles. An important next step will involve testing these model predictions using human experiments.

## 8. Generalizing TMS outcomes across modalities and cortical regions, and other future directions

Mean field models incorporating excitatory and inhibitory cortical populations have successfully described TMS plasticity effects observed in human motor cortex, using synaptic coupling strength as an analog for MEP amplitude. However, MEP formation is complicated, reflecting polysynaptic connections between cortical, corticospinal, and motor neurons (Ziemann et al., 2015). Indeed, changes in synaptic properties at the spinal level may contribute to MEP changes following rTMS (Perez et al., 2005; Quartarone et al., 2005). Future models should include spinal and motor neuron populations to more accurately capture the effect of TMS on the corticospinal system. Encouragingly, mean field models can also capture other TMS phenomena, such as short-interval cortical inhibition and intracortical facilitation following paired pulse paradigms (Wilson et al., 2014). More detailed models considering many discrete neurons in multiple layers are also able to capture these paired-pulse phenomena (Esser et al., 2005; Rusu et al., 2014), but at the cost of an increase in model complexity.

If a model of a neural system is a good approximation to the true biology, then it should be able to describe multiple phenomena with the same biophysically constrained parameters. Values for model parameters can be found either from experiments that measure them directly or by finding appropriate ranges that reproduce known effects. A good model must be able to generate predictions beyond the data used to constrain it. Mean field models, such as those described in this work, are capable of reproducing not just experimental results of TMS, but also a wide range of experimental phenomena, including oscillatory and evoked EEG activity (Rennie et al., 2002; Robinson et al., 2001), and slower hemodynamic oscillations measured with functional MRI (Steyn-Ross et al., 2009); these results can be used to further constrain model parameters.

TMS plasticity paradigms are often administered outside the motor system in cognitive and clinical applications where MEP measures are not possible and neuroimaging methods, such as EEG and fMRI, are increasingly used to assess how TMS alters neural activity in these non-motor regions (Sale et al., 2015; Thut and Pascual-Leone, 2010). Important future work will be to assess whether population models are capable of capturing TMS-induced changes in oscillations, event-related potentials, and hemodynamic oscillations.

Recent plasticity modeling with neural masses (Fung and Robinson, 2014; Wilson et al., 2016) has concentrated on the cortex alone. However, the thalamocortical system (Fig. 1C) is important for EEG dynamics, due to resonances in the thalamocortical feedback loop. Plasticity modeling may therefore be expanded to incorporate such dynamics in the future. As such, mean field models offer a unique opportunity to better understand the neural mechanisms driving TMS-induced changes in local cortical circuits outside of the motor system, and the impact of these changes across broader neural networks.

There are many unresolved factors regarding the interaction of magnetic pulses with the brain. The electric field induced in the brain by a TMS pulse can readily be modeled with finite element models with varying degrees of complexity, from simple geometries (Tang et al., 2016), through to folded human brain geometries extracted from MRI (Opitz et al., 2013) with the simNIBS tool (<u>www.simnibs.de</u>). In simple geometries, analytical calculations may be sufficient (Pashut et al., 2014), although how this field stimulates neurons is not yet fully known. This is of critical importance for modeling and interpreting TMS effects. A simple assumption is that the electric field gradient along a long axon triggers an action potential through buildup of transmembrane voltage (Roth and Basser, 1990). More detailed approaches use compartmental models for evaluation of the effect of the electric field on neural geometries and tracts (De Geeter et al., 2012; Pashut et al., 2011); these could be easily incorporated into a comprehensive modeling architecture, such as that outlined in Figure 6.

**Figure 6.**
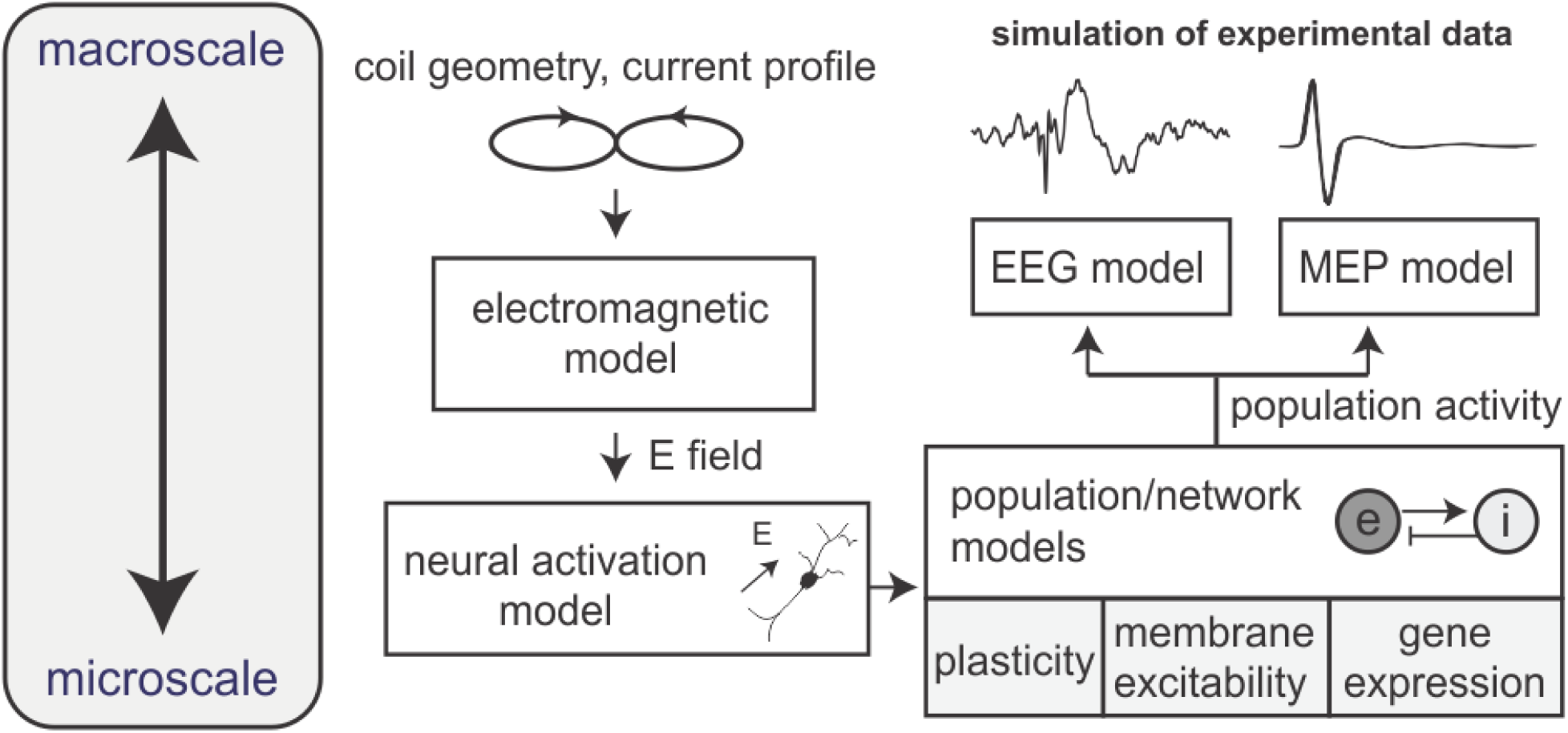
*The stages of a comprehensive modeling architecture. Stimulation with a given coil and current profile induces an electric field, which can be determined with an electromagnetic model. This electric field then causes changes in neural activation and behavior at a network level. This network activity then induces changes in the network, through a variety of possible mechanisms, including plasticity, changes in membrane excitability and gene expression. Finally, the altered activity can be mapped with appropriate models to measurable quantities such as EEG and MEP.*

Furthermore, while this work has discussed only plasticity, other mechanisms are likely to contribute to changes in neural activity following TMS. For instance, TMS-induced changes in membrane excitability (Pell et al., 2011) or the expression of biochemicals such as brain-derived neurotrophic factor (Gersner et al., 2011) could also influence plasticity and neural activity. Finally, this approach could be applied to other plasticity-inducing brain stimulation paradigms, such as transcranial direct current stimulation (Bestmann et al., 2015; Hämmerer et al., 2016). By incorporating these factors we aim to generate a powerful modeling framework in which brain stimulation can be systematically investigated and interpreted.

## 9. Summary

Understanding the mechanisms of TMS in humans is complicated by an interplay of different spatial and temporal scales. Neural plasticity and other possible drivers of TMS-induced effects occur at a microscopic scale, yet stimulation and measurement are made macroscopically. There is considerable variation in experimental results, since many of the key parameters, such as the initial cortical activity, are poorly controlled and differ between individuals. Moreover, the range of possible stimulating protocols is vast, but the range is mostly unexplored experimentally, with cTBS and iTBS protocols having dominated recent research. Mathematical modeling of the relevant physiology allows rapid evaluation of the effects of many paradigms and differing physiological states, and to explore the effects of microscopic changes on macroscopic outputs without the cost and time demands of experiment. We argue that the early modeling studies presented here hold great promise for providing a much needed theoretical framework with which to unify many diverse experimental findings and address many of the outstanding problems in the field.

Despite its transformative potential, TMS modeling is nascent. While the current models of TMS-induced plasticity have qualitatively captured several group-level findings following paradigms such as PAS and TBS, it remains to be seen whether these models can predict unobserved experimental findings, either at the group or individual level (e.g., the effect of changing the frequency of stimulation in TBS). An important next step is to design experiments that will directly test model predictions of unknown TMS parameters in order to assess the biophysical validity of these models. A goal for neural modeling is to design a general model capable of capturing a broad range of observed neural phenomena (e.g., neural oscillations, changes in brain states, event-related potentials). In order to test the generalizability of the plasticity models, future work will need to extend the current TMS plasticity models to incorporate spatial dynamics across cortical and subcortical regions (e.g., Fig. 1C). Continued development of models of TMS plasticity will enable closer integration between experimental and theoretical work to guide the field and unify diverse experimental findings in terms of underlying mechanisms.

## Acknowledgments

This work was supported by the Australian Research Council via the Center of Excellence for Integrative Brain Function (Grant CE140100007; PAR), Laureate Fellowship Grant (FL1401000225; PAR) and Future Fellowship (FT130100589; AF), and the National Health and Medical Research Council of Australia via a Project Grant (1104580; AF & NCR), and Fellowship (1072057; NCR).

